# Tech Note: Increased Complexity of Amplicon Libraries using Phased Primers

**DOI:** 10.1101/2025.08.26.672315

**Authors:** Franziska Bonath

## Abstract

Phased primers are used during the first PCR of amplicon library preparation in order to increase the complexity of the library. This increased complexity leads to improved sequencing quality even when sequencing with a 2×300 paired-end setup. We can show that libraries prepared using a phased primer approach are complex enough to reduce the amount of PhiX added to the sequencing reaction to that of other, complex libraries.

## Introduction

Short read amplicon sequencing is a cheap and fast way to determine both presence and absence as well as the sequence variation for a specific short sequence of interest in a variety of samples. During the amplicon library preparation the region of interest is amplified and sequences are added that are needed for the sequencing reaction on the flow cell. Additionally specific indexes are attached so that a large number of samples can be sequenced at the same time (Figure 1). In most cases this process is achieved in a 2-step PCR approach, in which the first PCR amplifies the region of interest and adds overhangs containing the priming site for the second PCR that adds sample specific indexes.

**Figure 1.**
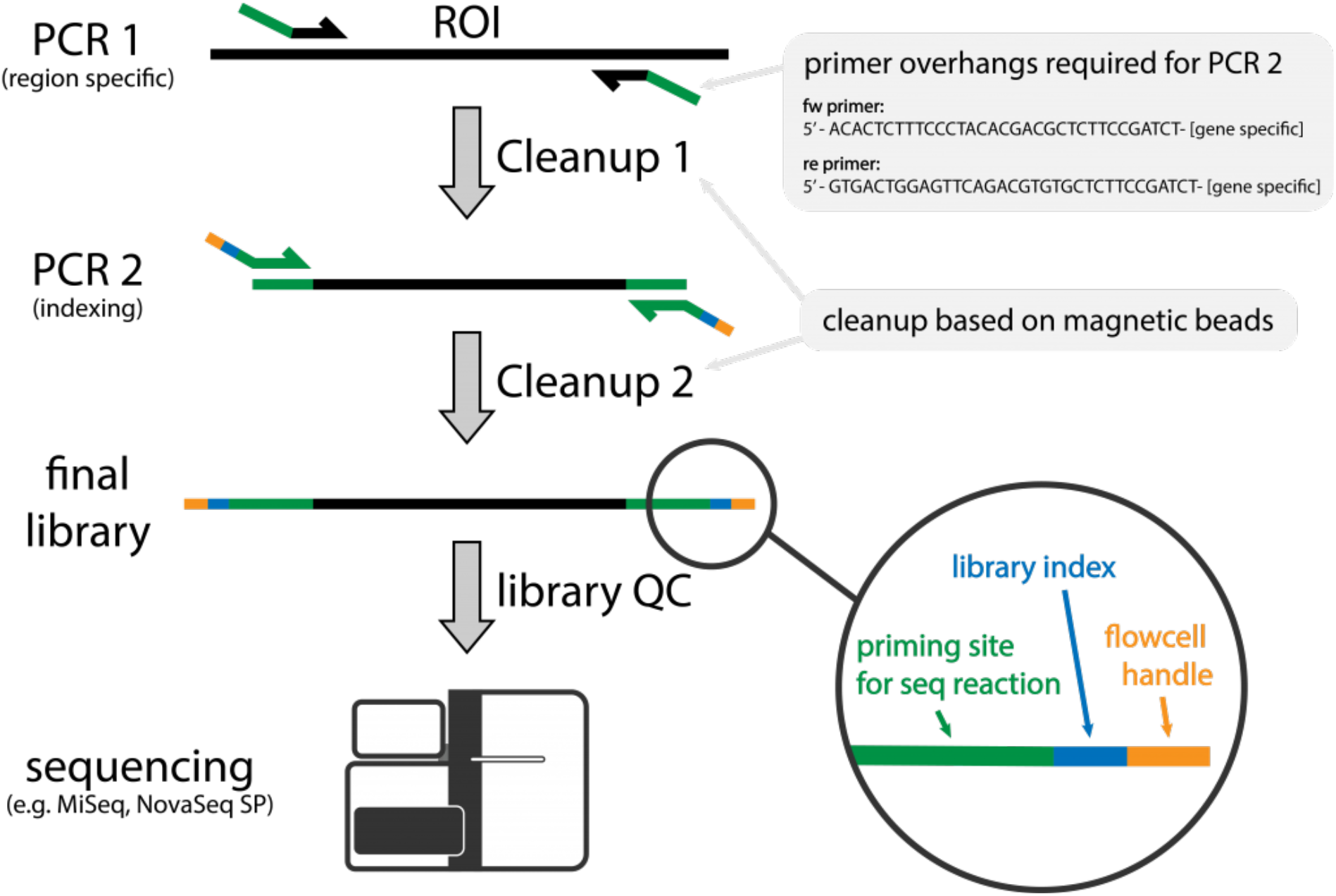
Schematic of Amplicon Library Preparation at NGI A region of interest (ROI) is amplified with region specific primers containing overhangs. During a second amplification sample specific indexes are added.

To date, the most commonly used sequencer for amplicon libraries is the Illumina MiSeq, since it can sequence up to 600 bp long sequences. However, the lack of sequence complexity, inherent to any amplicon library preparation, is a big concern when sequencing on Illumina machines. Every Illumina sequencing reaction begins with the generation of sequencing clusters: the local amplification of single molecules with flowcell bound primers. This is required to tether the molecule to the flowcell and to increase the signal. On a non-patterned flowcell, as can be found on MiSeq sequencers, these clusters are randomly distributed. As such, clusters can be close together or even overlap. During the early cycles of the sequencing reaction, the Illumina software identifies the different clusters and generates quality metrics based on the intensities found within them. If the sequence between overlapping clusters differs during these crucial cycles, the software can identify them as different clusters, however in less complex libraries, the likelihood of both molecules having the same sequence is high and hence the two clusters will be identified as one. Subsequently, the expected signal intensities for bases are incorrect and the base calling will be flawed leading to low sequencing quality.

For low complexity samples, Illumina currently advises to reduce the library concentration and to spike in 5-10% of a highly complex library, like the Illumina PhiX control. This will reduce the chance of clusters overlapping and increase the possibility of clusters with different sequences to be adjacent. Doing so, however, will also decrease the flowcell output since there will be a fewer clusters and some of the clusters will originate from the control library. Despite these measures, low complexity libraries still often suffer from lower sequencing quality, especially when sequencing longer fragments.

At the National Genomic Infrastructure we therefore use phased primers in the first PCR. This small change in our amplicon library preparation leads to highly complex amplicon libraries and subsequently improved sequencing quality even when sequencing with a 2×300 paired-end setup. We can show that libraries prepared using a phased primer approach are complex enough to reduce the amount of PhiX added to the sequencing reaction to that of other, complex libraries.

## Methods

### Overview Primer design for 16S amplicon library prep

To amplify the 16S region we use the 341F and 805R primer sequences as published in Hugerth et.al 2014. For non-phased library preparation, the indexing scheme of Hugerth et.al 2014 was implemented. Phased library preparation used the Adapterama indexing scheme adapted from Glen, Nilsen, et al., 2019 and Glenn, Pierson, et al., 2019. Further, we inserted up to 7 phasing nucleotides in both the forward and reverse primers of PCR1 as described in Wu et.al 2015 (for more information see the Results section) and used unique dual indexing in forward and reverse.

The primer sequences for PCR 1 are noted in Table 1, indexes used in the Andersson scheme in Table 2 and indexes in the Adapterama scheme in Table 3. All tables can be downloaded via the respective links at the end of the Tech Note.

### Bioinformatic Tools

We used RStudio version 1.1.463 with the shiny package to generate the R script for the shiny app. The code of the phased primer shiny app can be [downloaded from github](https://github.com/FranBonath/phased_primers_shiny/tree/main/phased_primers_shiny) or run online following this [link](https://fbonath.shinyapps.io/test_shiny/?_ga=2.236905921.1362636194.1610527973-1271399215.1610527973).

Analysis of library quality and sequence composition was performed using MultiQC via a standard sequencing QC pipeline at the National Genomic Infrastructure.

### Library Preparation Andersson Scheme

Library preparation for amplicon libraries using the Andersson scheme was performed on 257 samples based on the protocol published in Hugerth et.al 2014 with modifications. The PCR 1 reaction mixture contained 1 ng of input DNA, 10µl KAPA HiFi Hotstart 2x Master Mix (Roche, KK2602), 0.5µl BSA (20mg/ml,Thermo Fisher, B14), 1 µl of 10 µM AA_341F and 1 µl of 10 µM AA_805R. Nuclease free water was added to a reaction size of 21 µl. The PCR conditions for PCR1 in the Andersson scheme were as follows: 98 °C for 2 minutes, 20 cycles of 98 °C for 20 s, 54 °C for 20 s, 72 °C for 15 s and a final elongation at 72 C° for 2 min. Samples were purified using 21 µl of MagSI NGSPrep plus purification beads (TATAA, MDKT00010075) following the suppliers instructions and samples were eluted in 12 µl of Elution Buffer (Qiagen, 19086). 6 µl of the PCR 1 eluate was subjected to amplification with the following reaction setup: 10 µl KAPA HiFi Hotstart 2x Master Mix, 1 µl of 10 µM i7 index primer and 1 µl of 10 µM i5 index primer. Nuclease free water was added to a reaction size of 20 µl. The PCR conditions for PCR2 in the Andersson scheme were as follows: 98 °C for 2 minutes, 8 cycles of 98 °C for 20 s, 62 °C for 20 s, 72 °C for 15 s and a final elongation at 72 °C for 2 min. Samples were purified using 20 µl of MagSI NGSPrep plus purification beads.

### Library Preparation Adapterama Scheme

Library preparation was performed on 90 samples. Forward and reverse primer mix were prepared by combining eight phased primers each, at equimolar levels. For primer sequences see table 1. The PCR1 reaction mixture contained 1 ng of input DNA, 10 µl KAPA HiFi Hotstart 2x Master Mix, 0.5 µl BSA (20 mg/ml), 1 µl of 7.5 µM forward primer mix and 1 µl of 7.5 µM reverse primer mix. Nuclease free water was added to a reaction size of 21 µl. The PCR conditions for PCR1 in the Adapterama scheme were as follows: 98 °C for 2 min, 20 cycles of 98 °C for 20 s, 54 °C for 20 s, 72 °C for 15 s and a final elongation at 72 °C for 2 min. Samples were purified using 21 µl of MagSI NGSPrep plus purification beads following the suppliers instructions and samples were eluted in 12 µl of Elution Buffer. 6 µl of the PCR1 eluate was subjected to amplification with the following reaction setup: 10 µl KAPA HiFi Hotstart 2x Master Mix, 1 µl of 5 µM Adapterama i7 index primer and 1 µl of 5 µM Adapterama i5 index primer. For primer sequences see table 3. Nuclease free water was added to a reaction size of 20 µl. The PCR conditions for PCR2 in the Adapterama scheme were as follows: 98 °C for 2 min, 8 cycles of 98 °C for 20 s, 55 °C for 30 s, 72 °C for 30 s and a final elongation at 72 °C for 2 min. Samples were purified using 20 µl of MagSI NGSPrep plus purification beads.

### Sequencing

Libraries were pooled at equimolar levels and the molarity determined by qPCR using the Illumina library Quantification kit from Roche (KK4824). Pools were run on either MiSeq V3 2×300 with a loading concentration of 10 pM (non-phased) and 11 pM (phased) or on MiSeq Nano 2×250 with a loading concentration of 8 pM. PhiX spike in was 10% unless otherwise indicated. The same library pool of phased libraries was used for all sequencing runs described.

## Results

### Design of phased primers for 16S amplicon sequencing

A standard 16S library requires on average about 100,000 sequencing reads for a general estimation of the composition of a bacterial community. As such, the by far most popular sequencer for 16S projects is the MiSeq machine with an output of about 18 Million reads, allowing to multiplex 180 samples per sequencing run if the 100,000 reads per library has to be achieved. The MiSeq V3 sequencing kit is also the only available Illumina kit that can sequence 300 cycles in paired end sequencing (a total of 600 sequenced bases per read).

On MiSeq machines, information from cycles 4-7 of read 1 are used to identify the position of the sequencing clusters and in the first 25 cycles the metrics for base intensities, phasing and prephasing are determined. Hence, the complexity of the sequencing pool in the first 25 cycles is crucial for the quality of the whole sequencing run. Unfortunately, the sequence of amplicon libraries prepared using a two-step PCR approach is determined solely by the primer sequence used during PCR 1 and usually contains only few degenerate bases. Therefore, the most important region of the library in terms of complexity is by design the least complex.

The 341F and 805R primers are common sequences used to determine the bacterial composition of environmental samples, for example from air, soil or marine origins. The 341F primer contains only two degenerate nucleotides at position 9 and 12. We designed 8 forward and 8 reverse primers with 0-7 additional nucleotides positioned between the priming site for the sequencing primer and the region specific primer (Figure 2A) that will bind the ribosomal DNA of the gDNA sample. The nucleotides were chosen in order to maximize the complexity at every given position of the sequencing reaction (Figure 2B). These primers were used in a primer mastermix where all forward and reverse primers were added at equimolar levels during PCR 1. The PCR reaction was optimised for primer concentration and annealing temperature using *E*.*coli* gDNA as input. All conditions were tested with DNA input ranging between 0.1 ng and 5 ng (data not shown).

**Figure 2.**
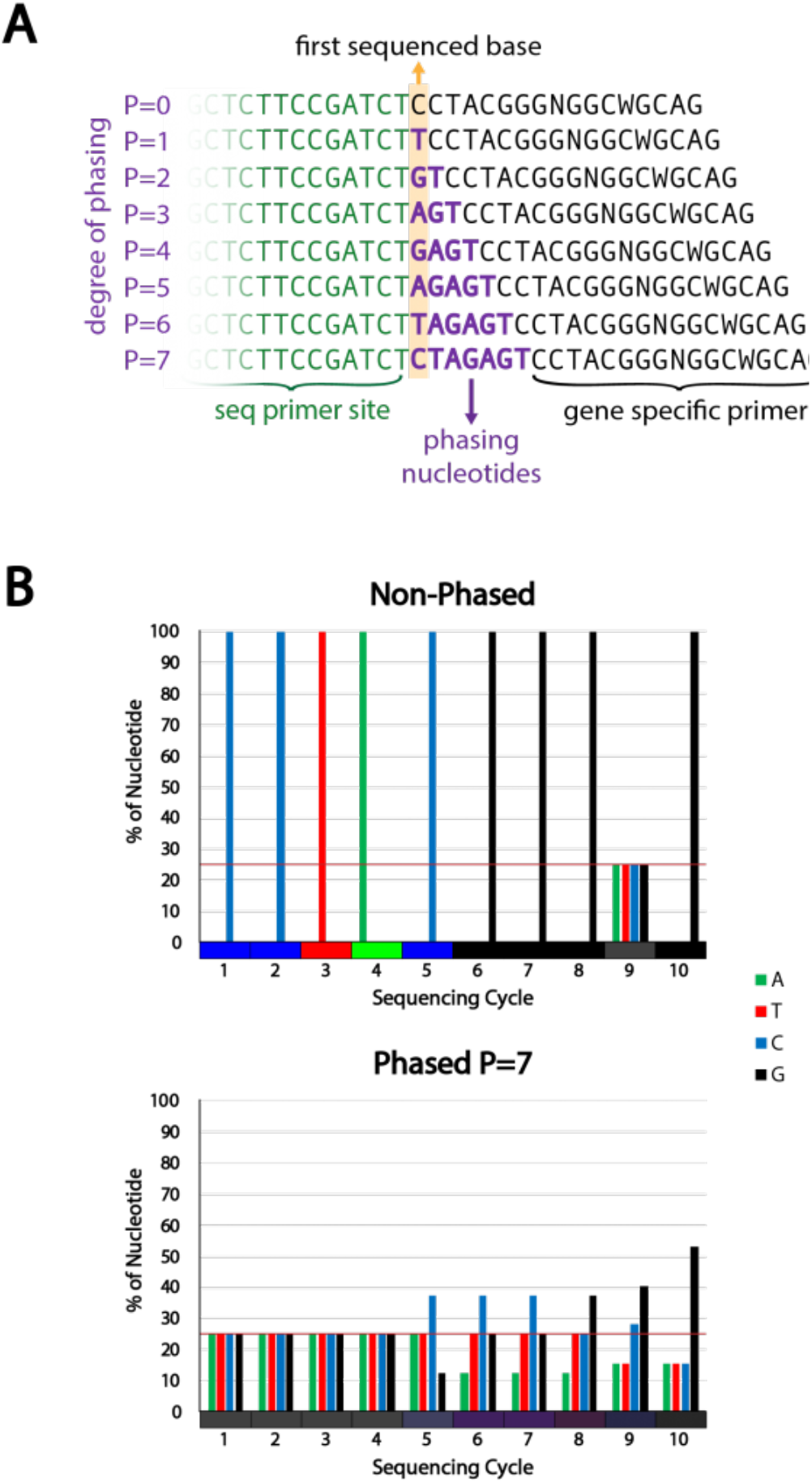
Phased primer strategy and design A) Strategy for phased primers. By addition of phasing nucleotides between the sequencing primer site and the gene specific primer sequence the nucleotide composition at different sequencing positions is increased in complexity. The degree of phasing determines the complexity. The phasing nucleotide sequence is chosen to induce maximum complexity at all positions. B) Example of the theoretical complexity change achieved with the 16S 341F primer using a phased approach with a phasing of 7 (bottom) compared to the non-phased primer (top). The complexity is also shown by mixing the different base colors in proportion to their share at each sequencing cycle position (color bar at the x-axis).

### Phased primers increase sequence complexity of libraries

We compared the effects of either the non-phased or phased library preparation on the complexity of samples submitted to NGI. As expected, non-phased libraries display near-digital nucleotide distributions at any cycle position that does not correspond to a degenerate nucleotide in the primer. Libraries that were prepared using the phased primer approach, in contrast, showed a high degree of nucleotide diversity, including in the crucial position 4-7 in read 1 (Figure 3).

**Figure 3.**
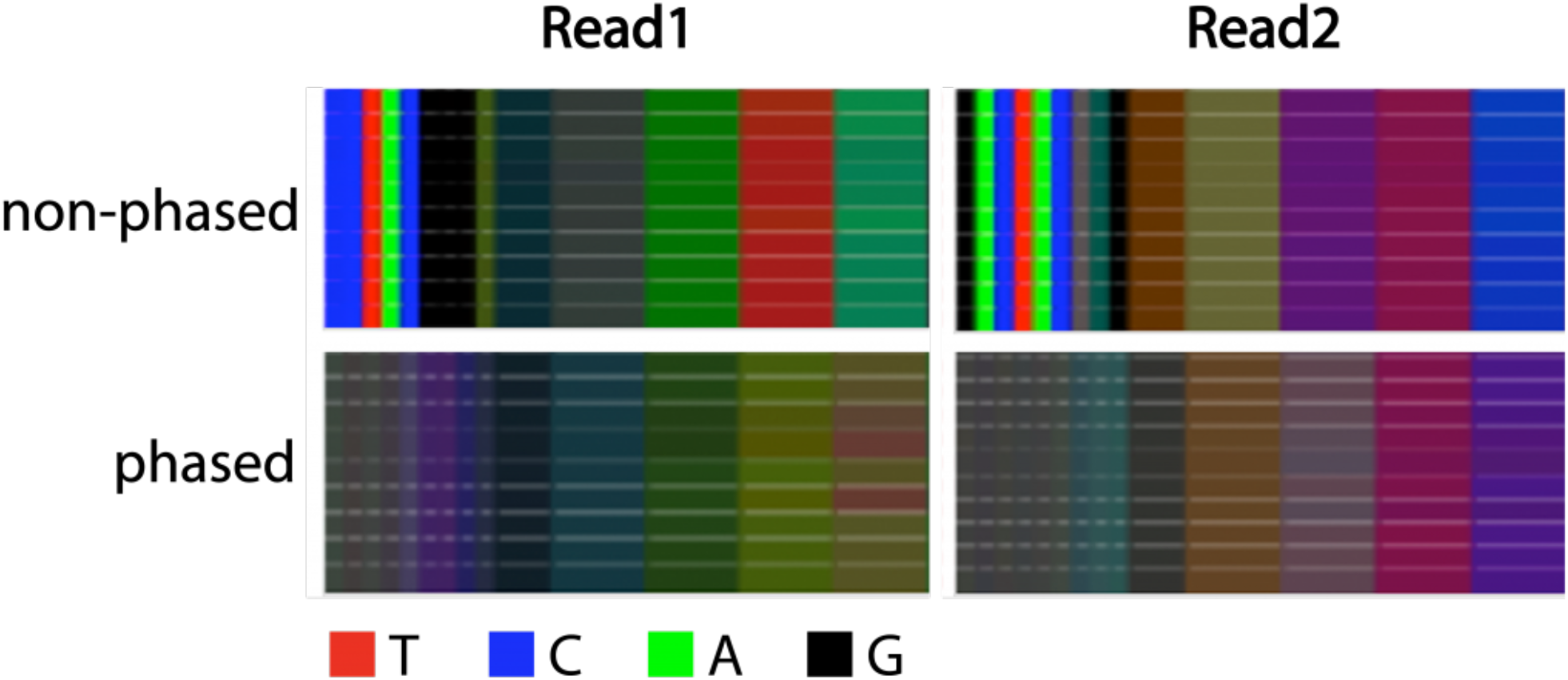
Complexity of libraries with and without phased primers Color bars displaying the proportion of the bases for the first 60 nucleotides of 10 libraries prepared with either phased or non-phased primers. Nucleotides 20-60 are shown in bins of 10 nucleotides. Read 1 derives from primer 341F and Read 2 from primer 805R.

### Phased primers increase the sequencing quality of libraries

Standard amplicon sequencing suffers from lowered sequencing quality despite the Illumina recommended addition of 10% PhiX. This is particularly pronounced for read 2 (Figure 4, non-phased library). We are using the Phred Score as a measure of quality, either for the quality at a specific sequencing cycle number over the length of the read (Figure 4A) or for the entirety of a given read (Figure 4B). As shown in Figure 4A, the read quality for read 1 is overall acceptable in non-phased libraries. Although individual libraries can show low qualities even in read 1. The quality of read 2 drops strongly starting from cycle number 200, falling below an average Phred Score of 30 at about cycle number 220 (Figure 4A). We would expect that a large amount of reads will have to be trimmed by 50 nucleotides or more.

**Figure 4.**
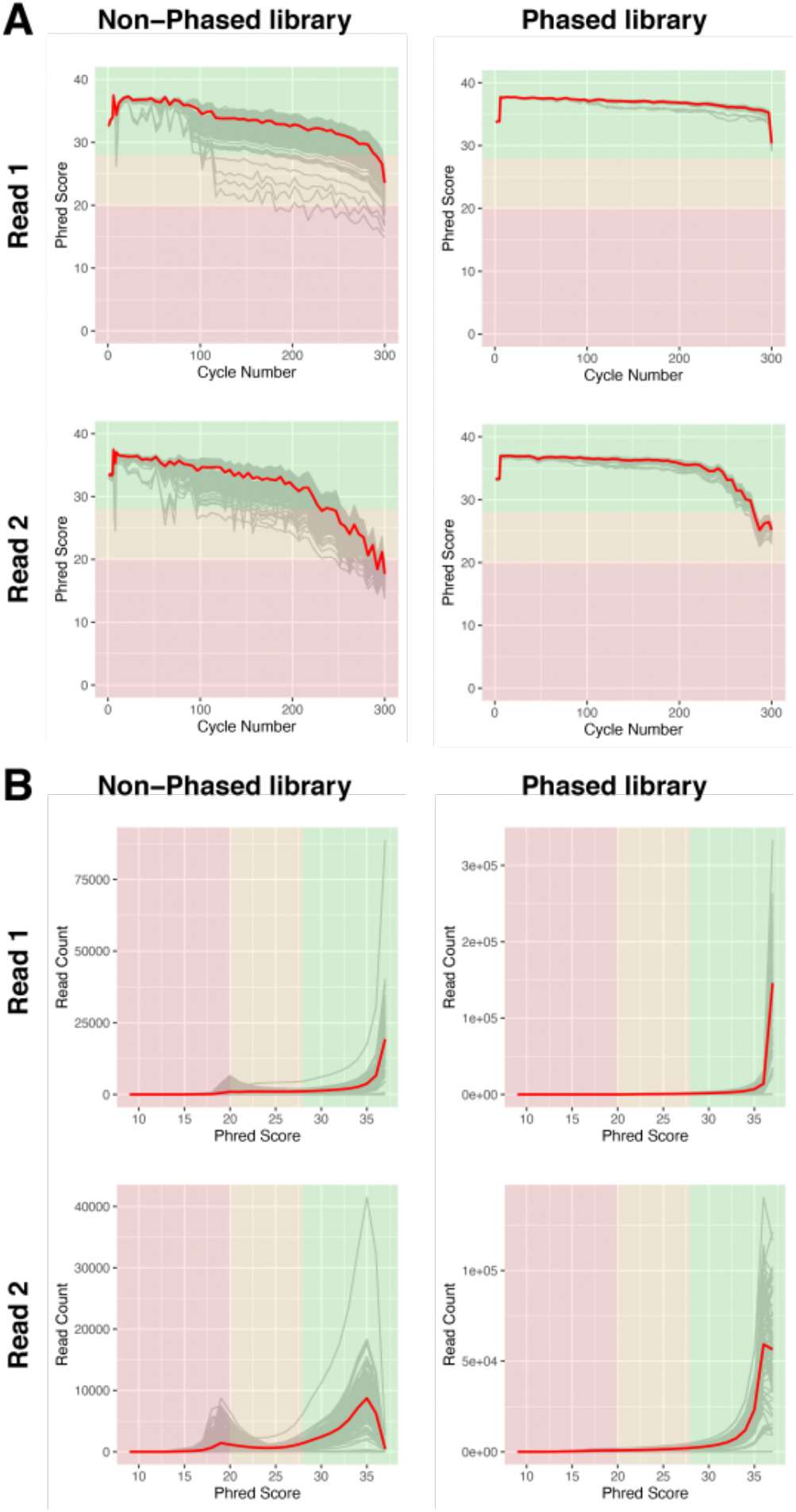
Sequencing quality of libraries with and without phased primers A) Base calling quality, measured by Phred Score, for each sequencing cycle in non-phased (left) or phased libraries (right). Grey lines indicate individual libraries, the red lines show the averages of all libraries of each graph. B) Base calling quality of reads for individual libraries (grey), measured by Phred score in non-phased (left) and phased libraries (right). The red lines show the averages of all libraries of each graph.

In stark contrast to non-phased libraries, the phased libraries show a very homogeneous read quality distribution that only falls moderately while progressing through read 1. As expected, read 2 shows a decline in read quality, however, the descent only starts at cycle 240 and drops on average below a Phred Score of 30 at about cycle 275 (Figure 4A).

This clear improvement is also apparent when inspecting Phred Scores by Sequence (Figure 4B). In non-phased libraries, there is an accumulation of low quality reads in all libraries, with a Phred Score of about 20, both in read 1 and read 2. These low quality reads are absent in phased libraries (Figure 4B).

Due to the low complexity of the libraries, we typically spike in 10% of PhiX libraries to mitigate at least some of the effects on sequencing quality. Since the phased libraries generated with the Adapterama indexing scheme appear to be highly complex we tested whether we can reduce the amount of PhiX to our standard input of 1% without compromising sequencing quality. To have the best possible comparison we used the same library pool run on the MiSeq V3 2×300 and resequenced them on MiSeq Nano 2×250 with either 10% PhiX or 1% PhiX. As shown in Figure 5, the pool with 1% spiked-in PhiX did perform comparably to 10% spiked-in PhiX in both, average quality over the course of the sequencing cycles (Figure 5A) as well as average read quality (Figure 5B).

**Figure 5.**
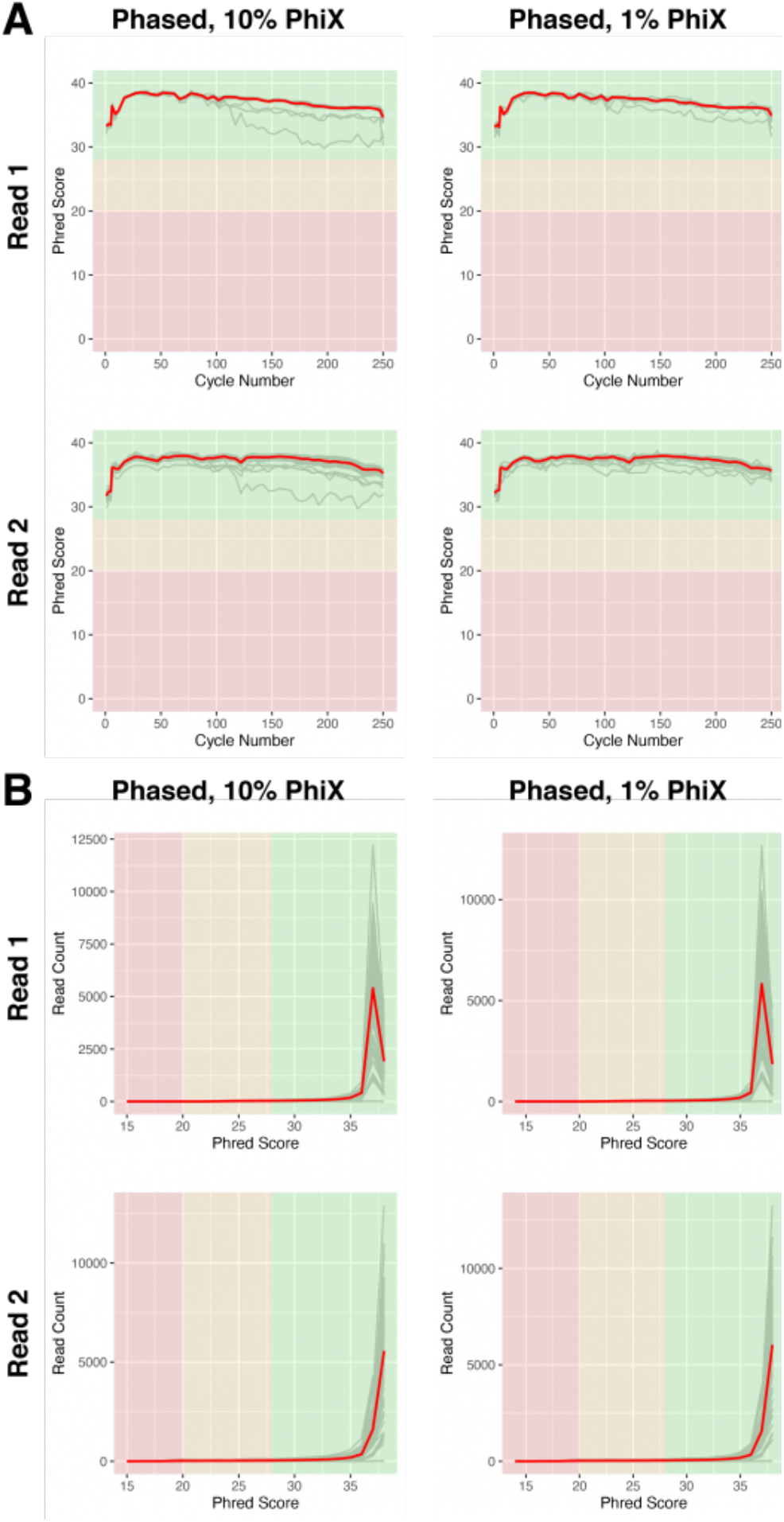
Libraries with phased primers do not require increased spike-in of PhiX A) Base calling quality, measured by Phred Score, for each sequencing cycle in phased libraries sequenced with 10% (left) or 1% (right) PhiX spike-ins. Grey lines indicate individual libraries, the red lines show the averages of all libraries of each graph. B) Base calling quality of reads for individual libraries (grey), measured by Phred score in phased libraries sequenced with 10% (left) or 1% (right) PhiX spike-ins. The red lines show the averages of all libraries of each graph.

### Simplified generation of phased primers

Since there is a clear improvement in sequencing quality when using phased primers, we want to aid researchers in their primer design. Therefore we created a web based tool which will automatically calculate the best nucleotide sequence for any given amount of phasing. The input requirements are the primer sequence to amplify the region of interest (Figure 6, (1)) as well as the overhang sequence that is used to prime for the second PCR (Figure 6, (2)). The user will be able to interactively change the degree of phasing (Figure 6, (3)) and the tool will return a graphic representation of the nucleotide distribution during the sequencing cycles 1-12 (Figure 6, (4)) and will display the primer sequences of all phased primers (Figure 6, (5)). The tool is using the IUPAC nomenclature for nucleotides and can also be applied to degenerate primers as long as it is using DNA bases only.

**Figure 6.**
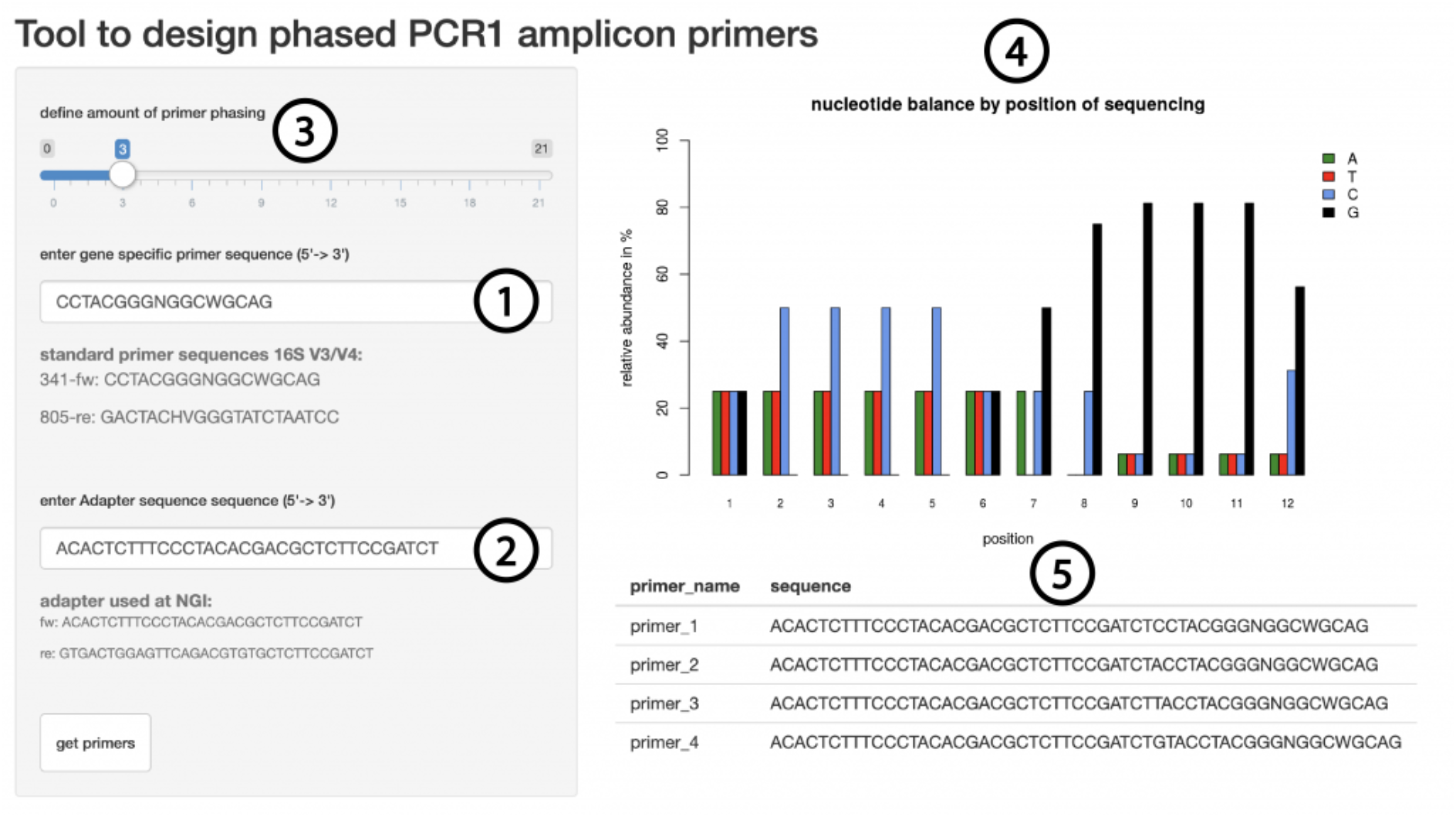
Shiny app to generate phased primers Users can enter the gene specific sequence (1), the overhang sequence (2) and a level of phasing (3) and will receive a graphic representation of the base complexity during the first 12 sequencing cycles (4) as well as the primer sequences required (5).

## Conclusions

Sequencing of low complexity libraries like amplicon libraries still poses a challenge to the current Illumina sequencing system. We evaluated the quality of sequencing using the MiSeq V3 2×300 sequencing kit, a popular choice among users with amplicon libraries. Our data shows that low complexity leads to decreased quality even when following the Illumina guidelines for sequencing of low complex libraries. Increasing the complexity of libraries by implementing phased primers during the library preparation increased the overall quality of the data leading to more reads that can be used in downstream analysis and less nucleotides that will need to be trimmed from each read due to poor quality.

Libraries prepared with phased primers show enough complexity in order to reduce the amount of PhiX spike ins to the levels of complex libraries. This will further increase the amount of usable reads.

Amplicon sequencing has a broad range of applications that reach far beyond the use in 16S analysis of bacterial communities. The phasing as we suggest here is dependent on the specific primer used to amplify a region of interest and therefore needs to be determined for every new set of primers. Doing so manually can be very time intensive. The tool we present here is a quick and easy way to identify the best degree of phasing for any sequence and will immediately return the primer sequences required. We hope this tool will reduce the obstacles in designing amplicon primers with phased primers.

## Supporting information

Table 1 | Primer sequences for 16S V3/V4 amplicon library preparation

Table 2 | Index sequences Andersson Scheme

Table 3 | Index sequences for Adapterama scheme (taken from Nextera UD, Illumina)

